## Supplemental fig. 1 - 9 for "Meiosis-specific decoupling of the pericentromere from the kinetochore"

**Extended Data Fig. 1**

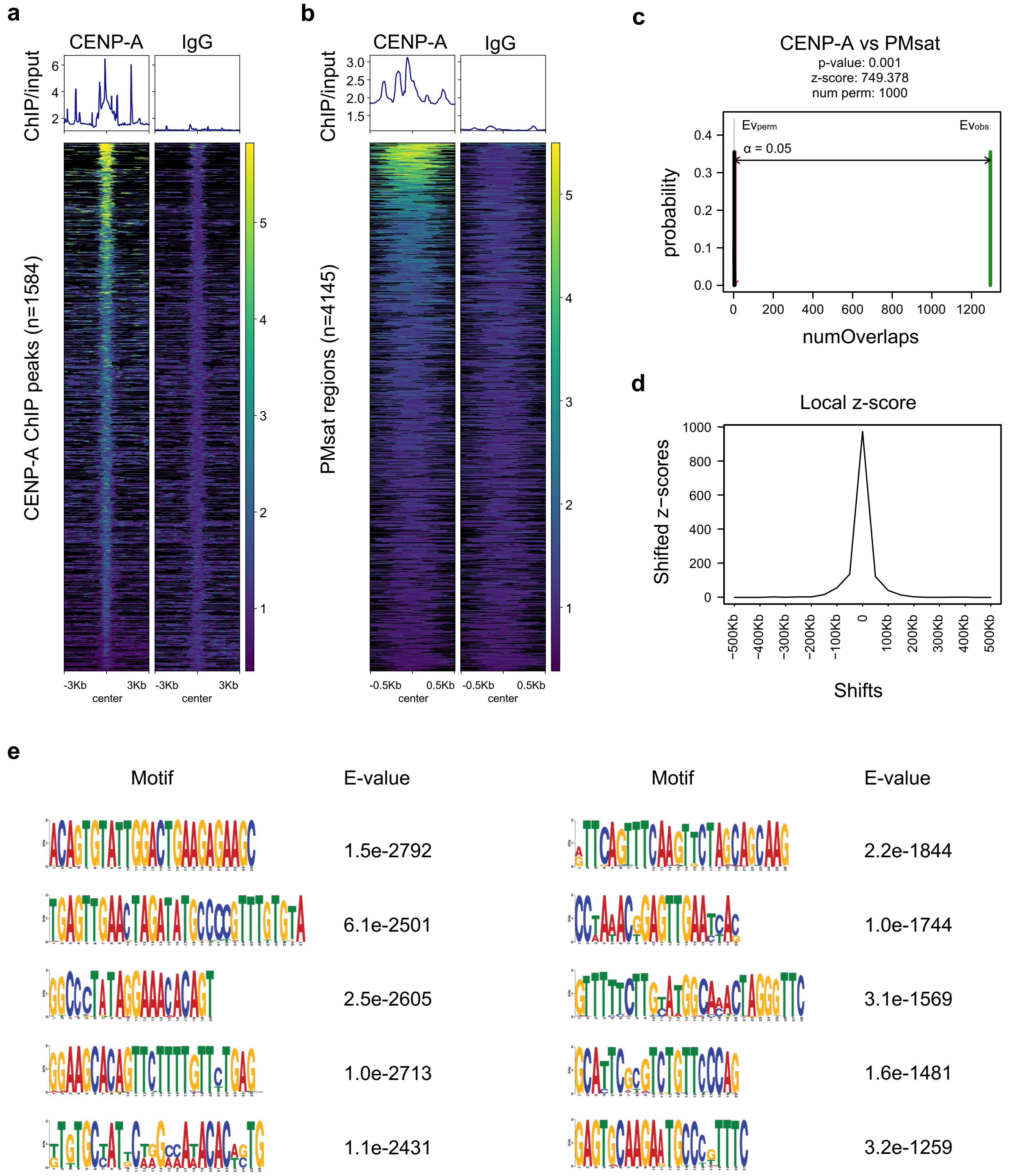


AT**ACAGTGTATTGGACTGAAGAGAAG**CTCCTGTTTTACTtAGTTTCCAATAC**TgAGTTGAACTAGATATGCCCCGTTTGTGTAGGAAGCACA**
**GTTCTTTTGTTCTGAG**CTAGCTCACTGTACT**GGCCCTATAGGAAACACAGT**A**GAGTGCAAGaGTGCCTGTTTC**TCAAAAGCAGAGT**GTTTTT**
**CTTgTAAGGCCGACTGGGGTTC**AATACAAcTT**CCTAAACGGAGTTGATTCACATTGAGTTTCAaGTTCTAGCATTAAG**aGTTAATTCAGGGT
AAGAACACCACTA**tTGTGCTATTCTGGCCATAAACAGTG**CAAATA**GCACTCGCGTCTGTTCCCAG**CAA

**Extended Data Fig. 2**

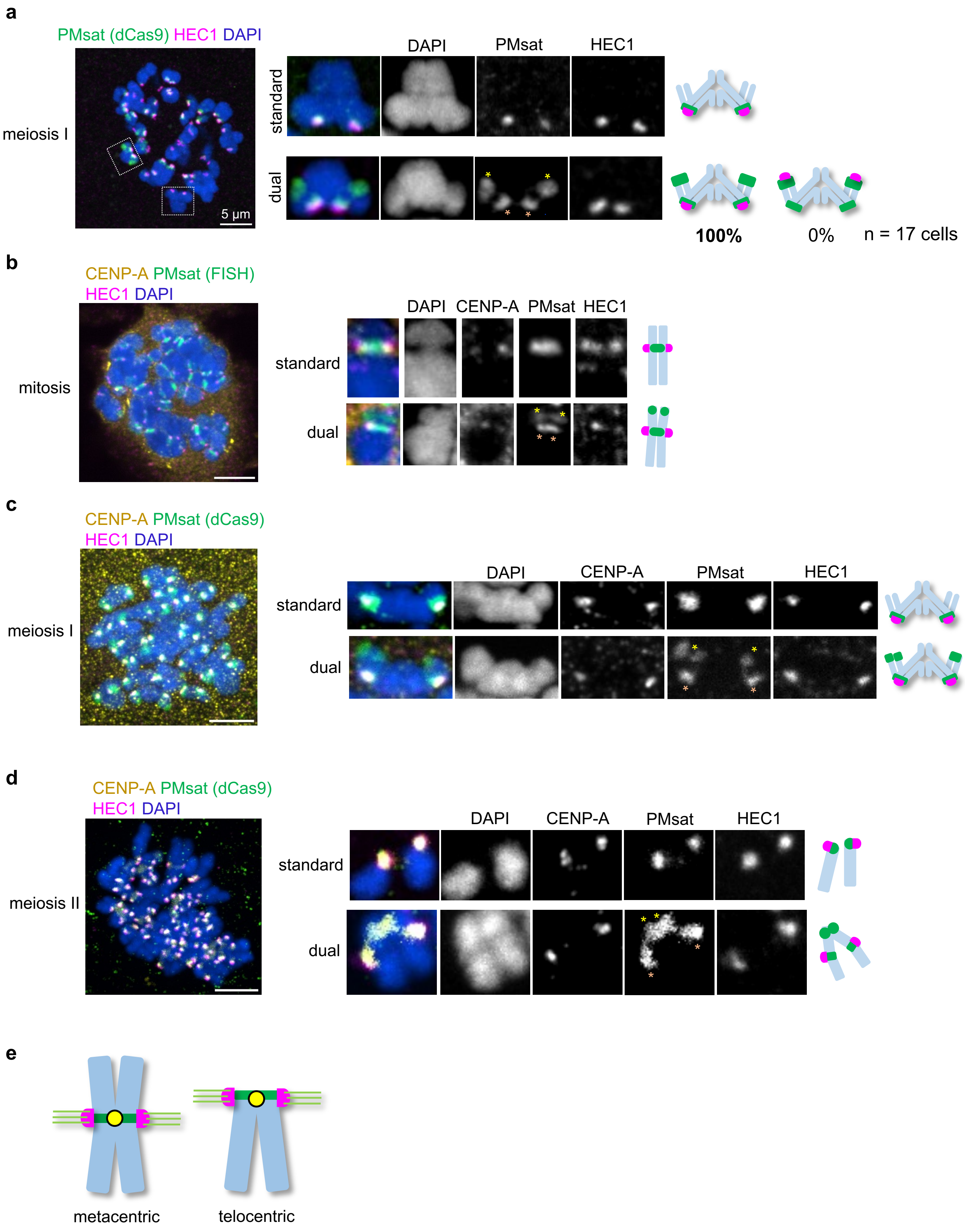

**Extended Data Fig. 3**

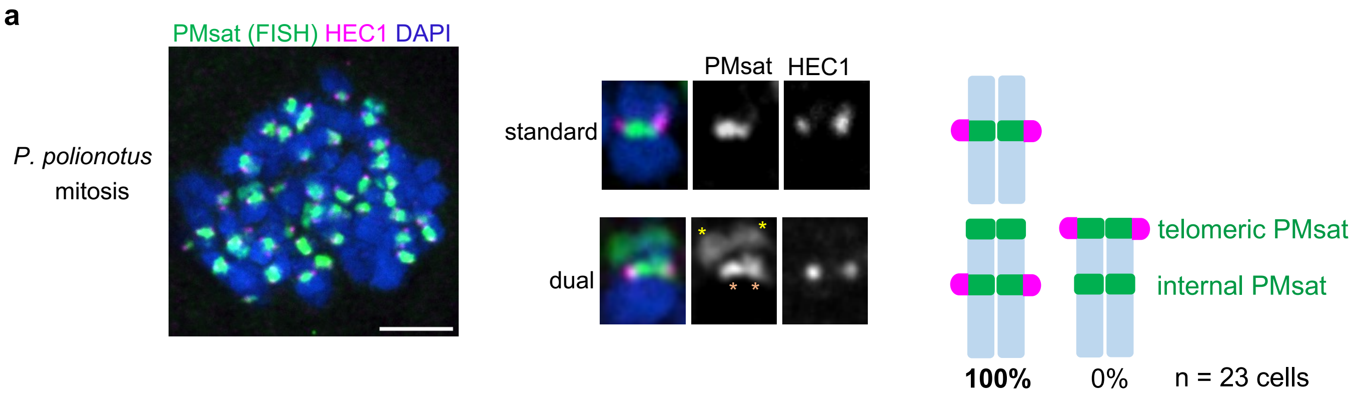


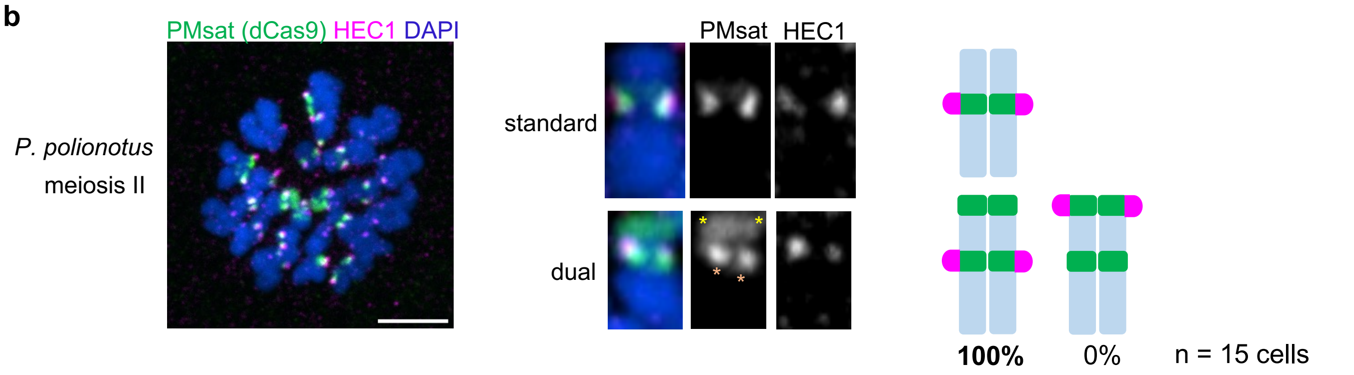

**Extended Data Fig. 4**

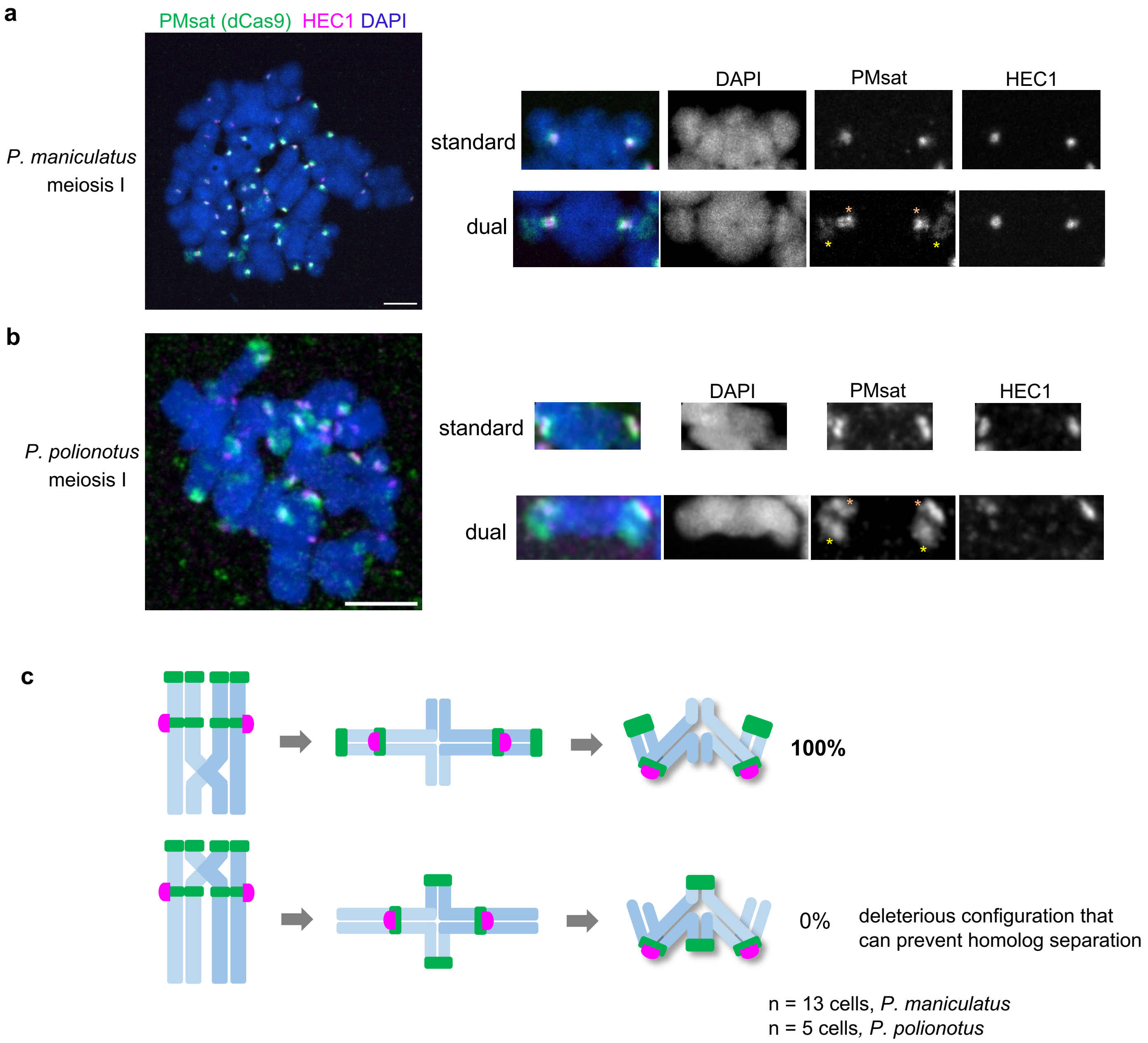

**Extended Data Fig. 5**

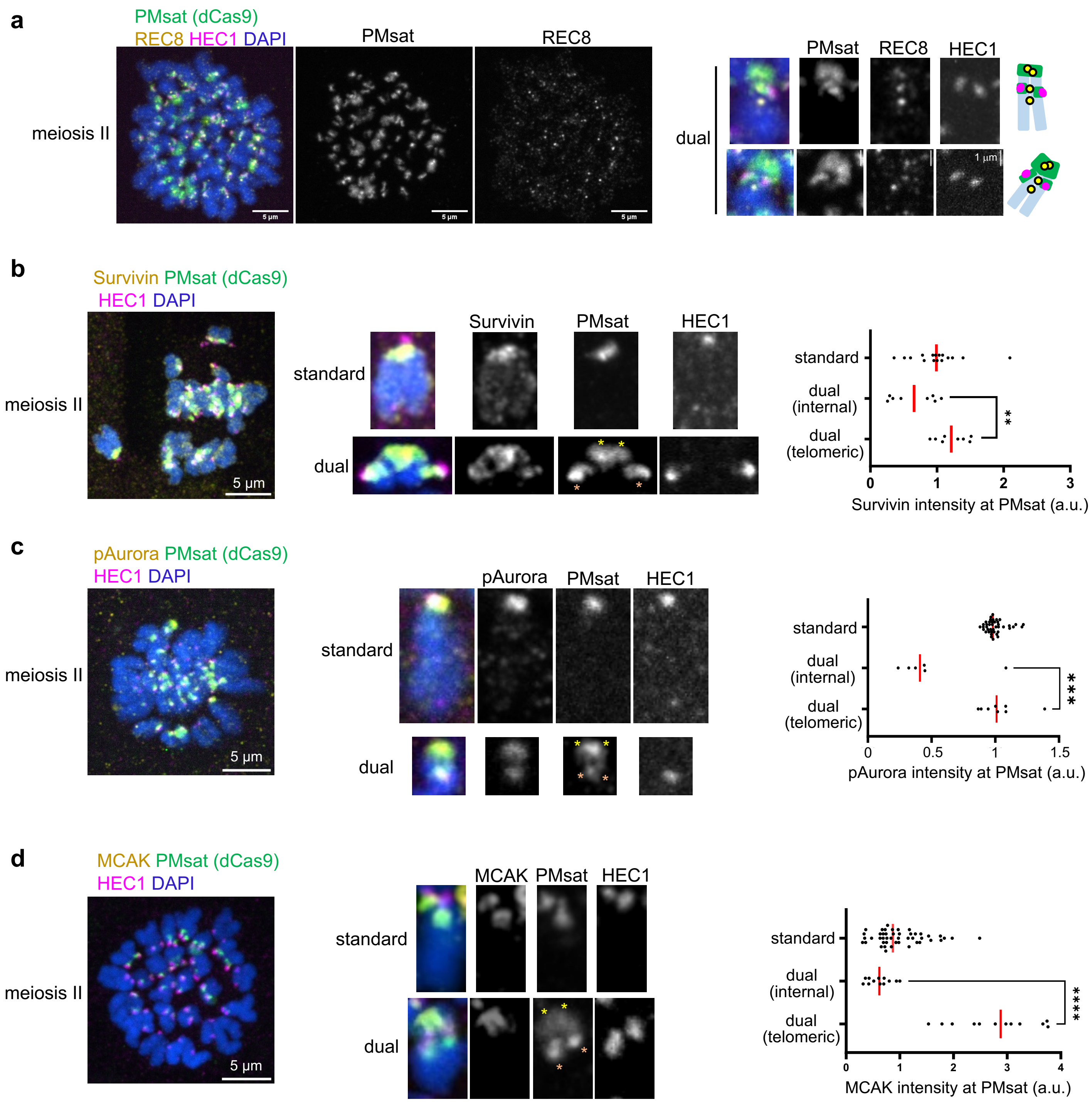

**Extended Data Fig. 6**

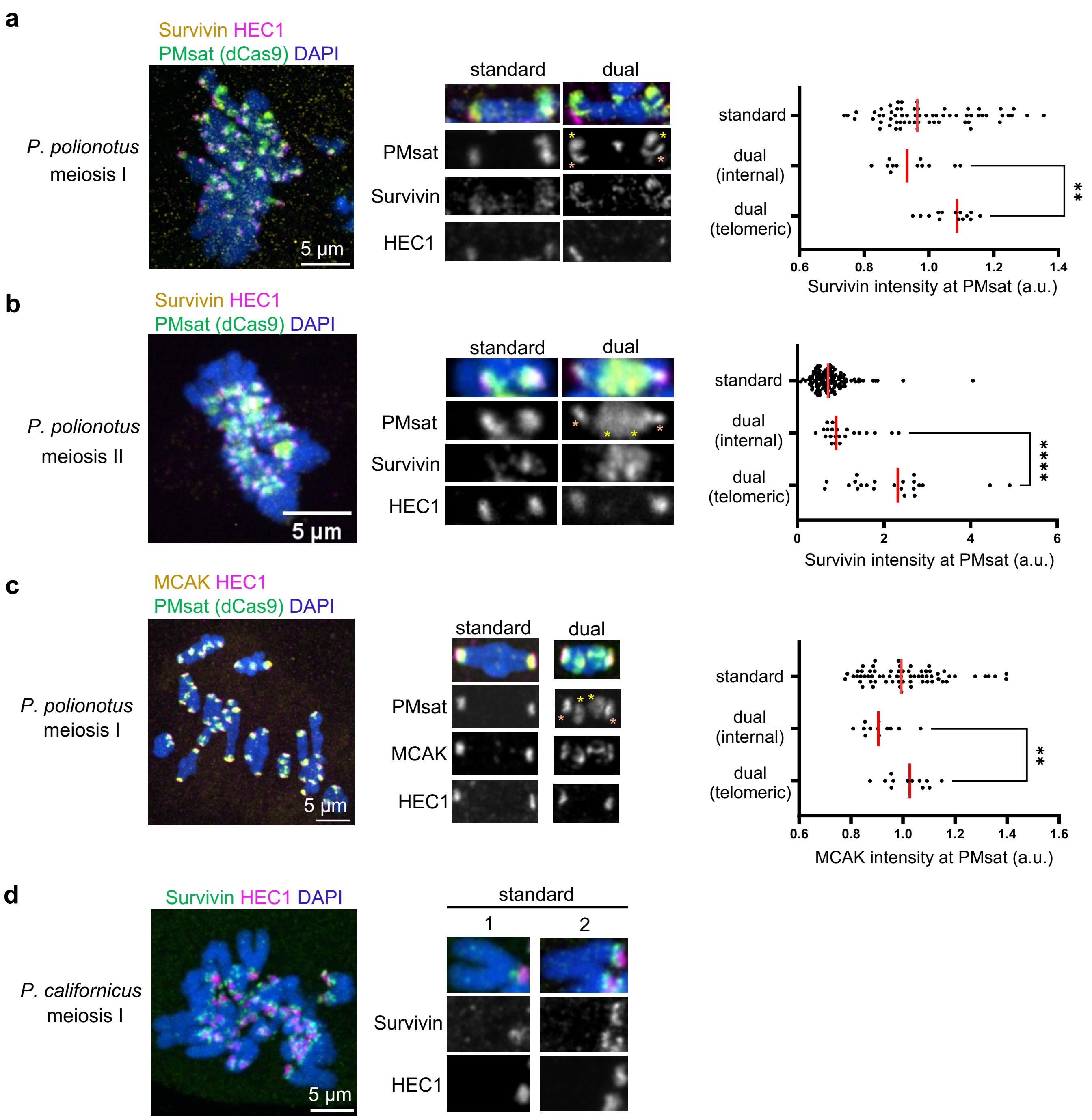

**Extended Data Fig. 7**

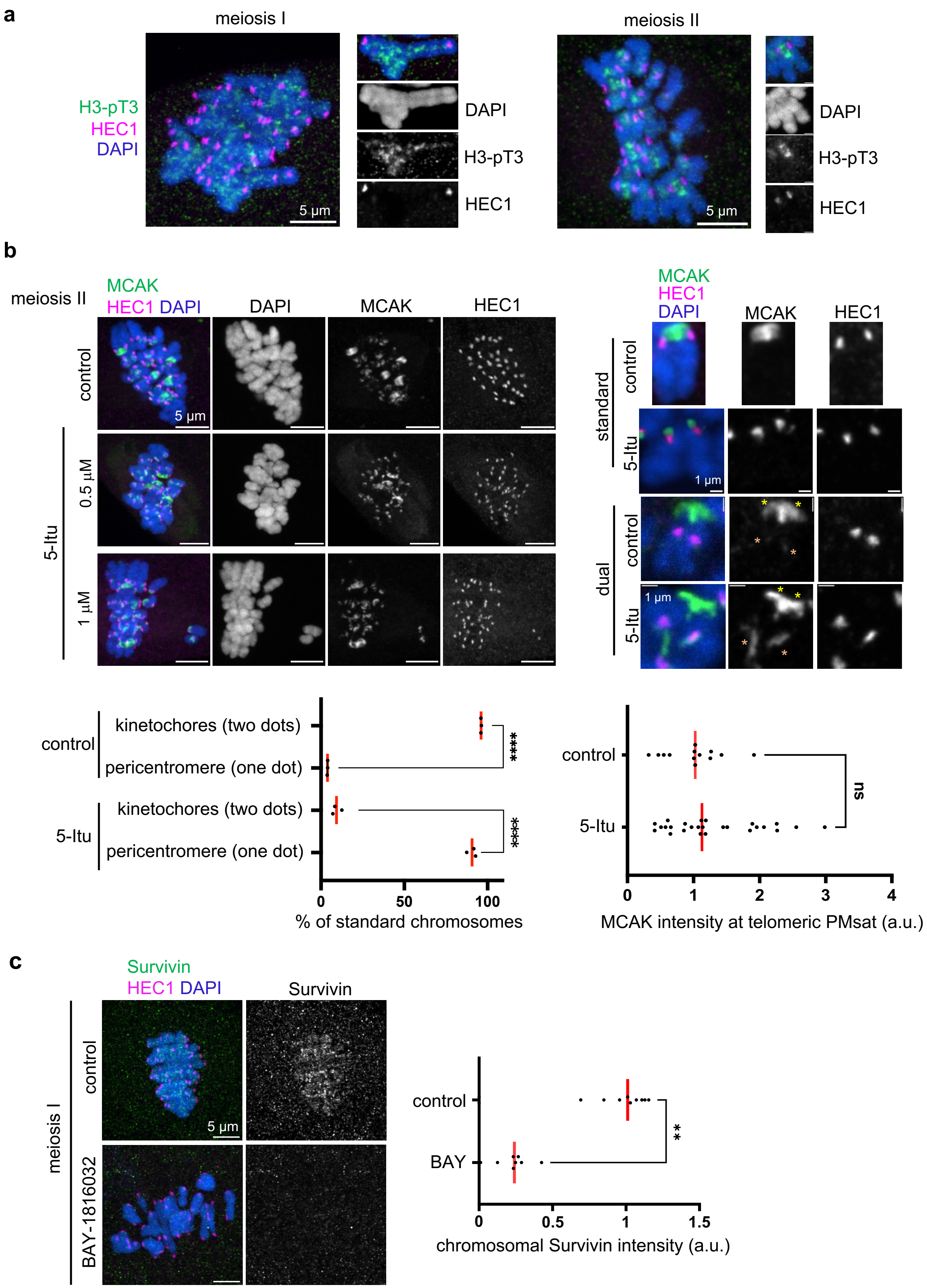

### Extended Data Fig. 8

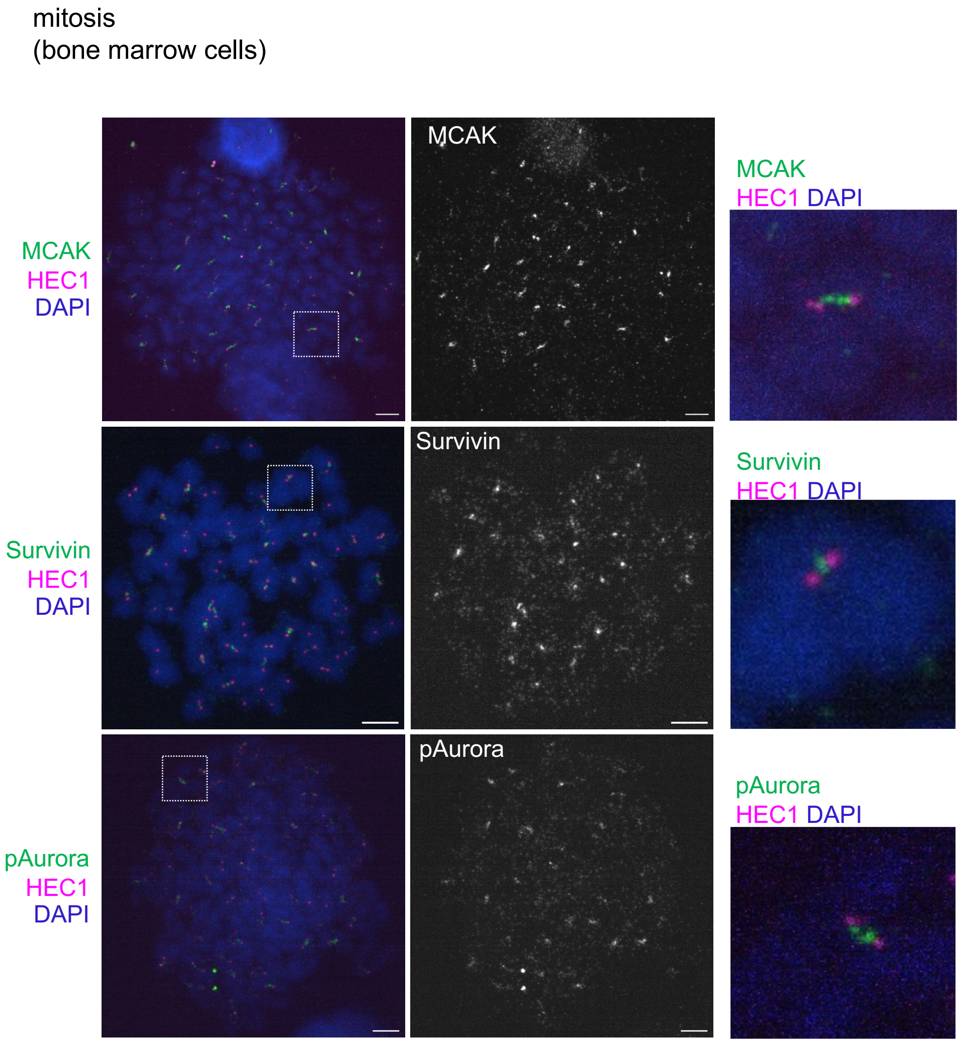

**Extended Data Fig. 9**

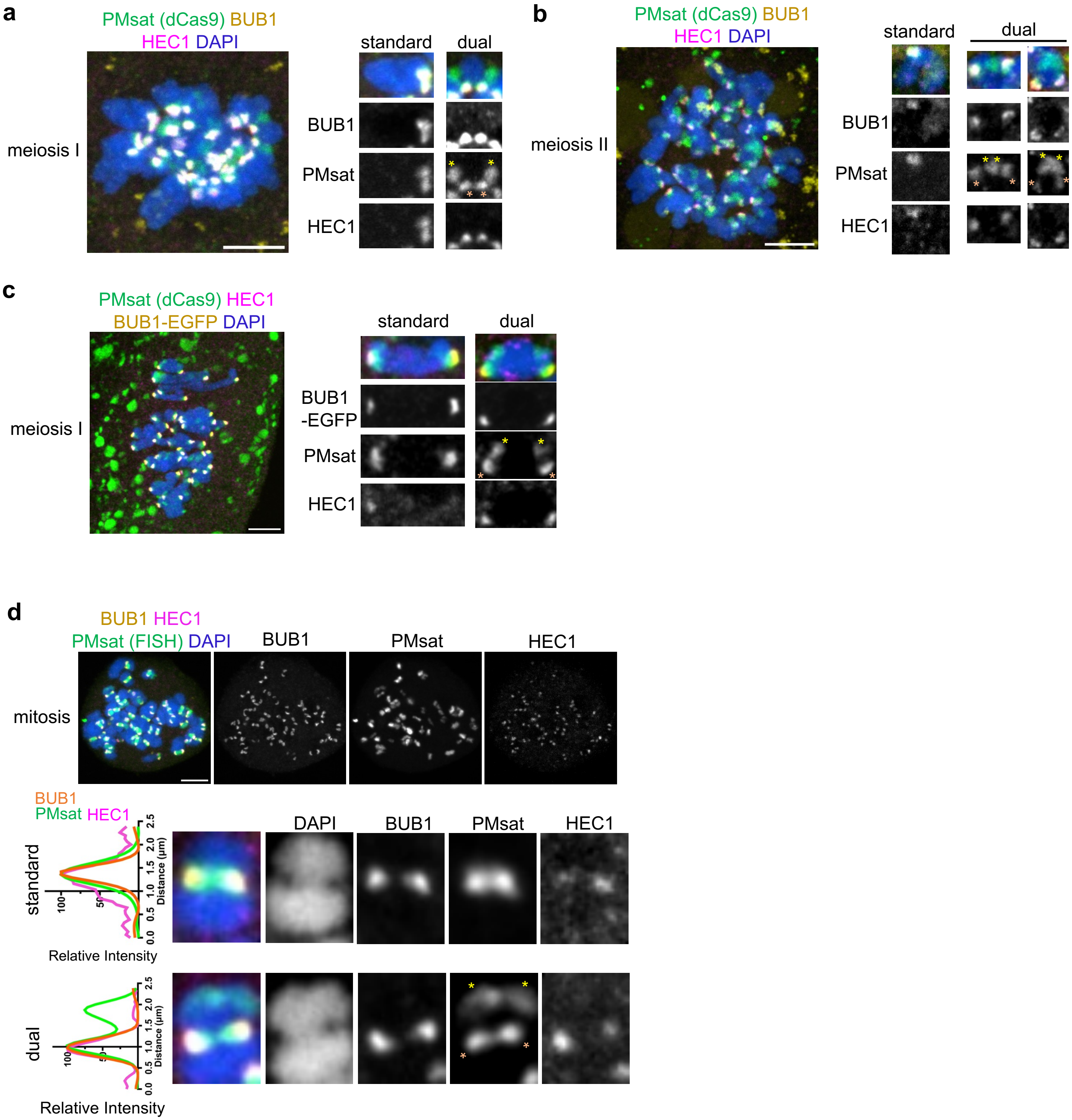
