## Supplemental Table 1 for "Meiosis-specific decoupling of the pericentromere from the kinetochore"

#### Main Figures

| | | experiment repeats | Number of chromosomes | Number of Cells |
|---|---|---|---|---|
| Fig 1c | mitosis | 3 | | 24 |
| Fig 1d | meiosis II | 5 | | 56 |
| Fig 2a | mitosis | 3 | | 13 |
| | meiosis II | 11 | | 46 |
| Fig 2b | meiosis I | 3 | | 14 |
| | meiosis II | 3 | | 9 |
| Fig 2c | IgG | 3 | | 23 |
| | REC8 | | | 15 |
| Fig 3a | | 3 | 152 | |
| Fig 3b | control | 4 | | 26 |
| | OA | | | 33 |
| Fig 4b | | 3 | 32 | |
| Fig 4c | | 3 | 210 | |
| Fig 4d | | 3 | 87 | |
| Fig 5a | control | 3 | | 16 |
| | 5-Itu | | | 15 |
| Fig 5b | control | 3 | 26 | |
| | 5-Itu | | 23 | |
| Fig 5c | control | 3 | | 13 |
| | BAY | | | 11 |
| Fig 5d | control | 3 | | 16 |
| | BAY | | | 10 |
| Fig 5e | control | 4 | 122 | |
| | BAY | | 141 | |
| Fig 6a | | 3 | | 13 |
| Fig 6b | | 3 | | 21 |
| Fig 6c | | 3 | | 11 |

### Extended Data Figures

| | | experiment repeats | Number of chromosomes | Number of Cells |
|---|---|---|---|---|
| Fig 2a | | 3 | | 17 |
| Fig 2b | | 3 | | 11 |
| Fig 2c | | 3 | | 12 |
| Fig 2d | | 3 | | 15 |
| Fig 3a | | 3 | | 23 |
| Fig 3b | | 3 | | 15 |
| Fig 4a | | 5 | | 13 |
| Fig 4b | | 3 | | 8 |
| Fig 5a | | additional example from the experiment in Fig. 2b | | |
| Fig 5b | | 3 | | 14 |
| Fig 5c | | 3 | | 16 |
| Fig 5d | | 3 | | 10 |
| Fig 6a | | 3 | 98 | |
| Fig 6b | | 3 | 156 | |
| Fig 6c | | 3 | 91 | |
| Fig 6d | | 3 | | 30 |
| Fig 7a | | 3 | | 16 |
| Fig 7b | control | 3 | 25 | |
| | 5-Itu | | 34 | |
| Fig 7c | control | 4 | | 13 |
| | BAY | | | 14 |
| Fig 8 | MCAK | 3 | | 7 |
| | Survivin | | | 9 |
| | pAurora | | | 9 |
| Fig 9a | | 3 | | 8 |
| Fig 9b | | 3 | | 27 |
| Fig 9c | | 3 | | 21 |
| Fig 9d | | 3 | | 10 |
